# Parallel diversifications of *Cremastosperma* and *Mosannona* (Annonaceae), tropical rainforest trees tracking Neogene upheaval of the South American continent

**DOI:** 10.1101/141127

**Authors:** Michael D. Pirie, Paul J. M. Maas, Rutger A. Wilschut, Heleen Melchers-Sharrott, Lars W. Chatrou

## Abstract

*This preprint has been reviewed and recommended by Peer Community In Evolutionary Biology (http://dx.doi.org/10.24072/pci.evolbiol.100033).* Much of the immense present day biological diversity of Neotropical rainforests originated from the Miocene onwards, a period of geological and ecological upheaval in South America. We assess the impact of the Andean orogeny, drainage of lake Pebas, and closure of the Panama Isthmus on two clades of trees (*Cremastosperma*, c. 31 spp.; and *Mosannona*, c. 14 spp.; both Annonaceae) found in humid forest distributed across the transition zones between the Andes and Western (lowland) Amazonia and between Central and South America. We inferred phylogenies based on c. 80% of recognised species of each clade using plastid and nuclear encoded sequence markers, revealing similar patterns of geographically restricted clades. Using molecular dating we showed that diversifications in the different areas occurred in parallel, with timing consistent with Andean vicariance and Central American geodispersal. In apparent contradiction of high dispersal abilities of rainforest trees, *Cremastosperma* clades within Amazonia are also geographically restricted, with a southern/montane clade that appears to have diversified along the foothills of the Andes sister to one of more northern/lowland species that diversified in a region once inundated by lake Pebas. Ecological niche modelling approaches show phylogenetically conserved niche differentiation, particularly within *Cremastosperma*. Niche similarity and recent common ancestry of Amazon and Guianan *Mosannona* species contrasts with dissimilar niches and more distant ancestry of Amazon, Venezuelan and Guianan species of *Cremastosperma* suggesting that this element of the similar patterns of disjunct distributions in the two genera is instead a biogeographic parallelism, with differing origins. The results provide further independent evidence for the importance of the Andean orogeny, the drainage of Lake Pebas, and the formation of links between South and Central America in the evolutionary history of Neotropical lowland rainforest trees.

## Introduction

The immense biological diversity of the Neotropics is the net result of diversification histories of numerous individual lineages (Antonelli and Sanmartín, 2011; Hughes et al., 2013; Pennington et al., 2015). Plants and animals encompassing a wide spectrum of forms, life histories, and ecological tolerances have diversified in ecosystems ranging from high alpine-like conditions of the Andean Páramo, through to seasonally dry tropical forests and the humid forests of lowland Amazonia (Hoorn and Wesselingh, 2009; Hughes et al., 2013; Luebert and Weigend, 2014; Pennington et al., 2010). The dynamic geological and ecological contexts of Neotropical species radiations shift both in space and through time (Graham, 2011; Hughes et al., 2013). Understanding the importance of different factors in driving the origins of biological diversity therefore requires approaches that directly compare biologically equivalent species radiations within the same geographical areas and evolutionary timescales with the ecological conditions that prevailed in those times and places (Fine et al., 2014; Koenen et al., 2015; Terra-Araujo et al., 2015).

Even across seemingly similar ecosystems and organisms, there are differences in levels of biodiversity within the Neotropics. For example, comparing across Neotropical rainforests, tree alpha-diversity peaks in the wetter, less seasonal part of Western Amazonia (Hoorn et al., 2010; Hoorn and Wesselingh, 2009). Correlation of this diversity with particular current conditions, such as climate and soils, may suggest a causal link in sustaining, and perhaps even driving, diversity (Kristiansen et al., 2012; Stropp et al., 2009). However, both species diversity and ecological conditions have changed dramatically in the Neotropics since the Oligocene (Hoorn et al., 2010; Hughes et al., 2013). Hoorn et al. (2010) reviewed evidence including from the microfossil record suggesting a c. 10 to 15% increase of plant diversity between c. 7 and 5 Ma. This was shortly after the Late Miocene draining of a wetland system in Western Amazonia, known as the Pebas system, or lake Pebas, which existed from c. 17 to c. 11 Ma (Hoorn et al., 2010). The Late Miocene draining of Western Amazonia was a direct result of the uplift that caused orogeny in the Andes (Shephard et al., 2010), with continuous discharge of weathered material from the rising Andes leading to the gradual eastwards expansion of terra firme forests. Hoorn et al. (2010) concluded that the establishment of terrestrial conditions in Western Amazonia was a possible prerequisite for the (rapid) diversification of the regional biota.

The Andean orogeny itself has been linked to diversification, also in lowland (rather than just montane or alpine) biota (Fine et al., 2014; Gentry, 1982; Luebert and Weigend, 2014; Sarkinen et al., 2007). The timeframe of the influence of the Andean orogeny on Neotropical vegetation may extend back to the Miocene, i.e. from c. 23.3 Ma onwards (Burnham and Graham, 1999). However, much of the uplift occurred in the late Miocene and Pliocene (Gregory-Wodzicki, 2000) with intense bursts of mountain building during the late middle Miocene (c. 12 Ma) and early Pliocene (c. 4.5 Ma) (Hoorn et al., 2010). A further geological influence, the closing of the isthmus at Panama, facilitating biotic interchange between North and South America, also occurred during the Pliocene (Hoorn et al., 2010) and may also have driven diversification. The Panama isthmus has long been assumed to have fully closed by 3.5 Ma. This date has been challenged recently by fossil data and molecular phylogenetic analyses that may indicate migration across that Panama isthmus dating back to the Early Miocene, c. 6-7 Ma (Bacon et al., 2013, 2015; Thacker, 2017; Winston et al., 2017). Finally, distribution shifts along the Andean elevational range during climatic changes in the Pleistocene (c. 1.8 Ma onwards) may also have driven diversification (Hooghiemstra and van der Hammen, 1998).

These events describe an explicit temporal framework for various plausible causes of diversification in Neotropical taxa. The means to test these hypotheses is presented by clades distributed across the transition zones between the Andes and Western (lowland) Amazonia and between Central and South America. Just such distribution patterns are observed in a number of “Andean-centered” (sensu Gentry, 1982) genera of Annonaceae, *Cremastosperma*, *Klarobelia Malmea* and *Mosannona*. These were the subject of analyses using phylogenetic inference and molecular dating techniques by Pirie et al. (2006), who concluded that their diversifications occurred during the timeframe of the Andean orogeny. Pirie et al. (2006) also identified clades within *Mosannona* endemic to the west and east of the Andes and estimated the age of their divergence to be c. 15-6 Ma. Central American representatives of the predominantly Asian Annonaceae tribe Miliuseae (Chaowasku et al., 2014; Ortiz-Rodriguez et al., 2016) are not found east of the Andes, which might also suggest that the Andes formed a barrier to dispersal prior to the closure of the Panama isthmus.

Even if diversifications occurred within the same timeframe, they were not necessarily driven by the same underlying factors. Further phylogenetic inference approaches would allow us to test whether Andean-centered distributions that originated in *Cremastosperma*, *Mosannona*, and other Annonaceae clades were indeed influenced by common biogeographic processes (such as vicariance caused by Andean uplift), and whether allopatric speciation has been an important underlying process. For example, by combining phylogeny and species distribution modeling in an African clade of Annonaceae, Couvreur et al. (2011b) could show that niche differences among closely related (but mostly not co-occurring) species were more similar than expected by chance. This was interpreted as suggesting allopatric speciation driven by landscape changes (Couvreur et al., 2011). Such analyses might also contribute to testing diversification scenarios in Andean-centered Neotropical Annonaceae. However, results presented in Pirie et al. (2006) were based on limited taxon sampling within the genera, and species level relationships were largely unresolved, with the geographic structure apparent within *Mosannona* not reflected by supported clades in the other genera, limiting the power of phylogenetic approaches in general.

In this paper, we focus on two clades of trees found in Neotropical humid forest. Species of *Cremastosperma* (c. 31 spp.; Pirie et al., 2005) and *Mosannona* (c. 14 spp.; Chatrou, 1998) (both Annonaceae) occur from lowland (i.e. up to 500 m) through pre-montane (500-1,500 m) and into lower elevation montane rainforest, predominantly in areas surrounding the Andes in South America but also extending north into Central America (Chatrou, 1997). No species occur on both sides of the Andean mountain chain, suggesting that the Andes represents a current barrier to dispersal. In general, few species of *Cremastosperma*, and none of *Mosannona*, co-occur, which may suggest diversification driven by allopatric speciation. Those species of *Cremastosperma* with overlapping distributions are mostly limited to northern Peru and Ecuador. Our aims are to reassess the timing and sequence of shifts in ancestral distributions in *Cremastosperma* and *Mosannona* to test the influence of the following factors on species diversification in the Neotropics. 1) The northern Andean orogeny, dividing western and eastern lineages; 2) the drainage of lake Pebas creating new habitat in western Amazonia; and 3) the closure of the Panama Isthmus allowing geodispersal of lineages into Central America. To this end, we use phylogenetic inference, molecular dating and niche modelling techniques with new datasets for *Cremastosperma* and *Mosannona* based on expanded sampling of taxa and DNA sequence markers.

## Materials and Methods

### Taxon sampling

This study largely utilised previously unpublished sequence data (partly used in Pirie et al., 2005b), as well as published sequences (Mols et al., 2004; Pirie et al., 2005a, 2006, 2007; Richardson et al., 2004) (see Appendix 1). Datasets were constructed for *Cremastosperma* and *Mosannona* in two separate studies employing different outgroup sampling. For the *Cremastosperma* dataset, ten Malmeoideae outgroup taxa were selected: seven of tribe Malmeeae, including two accessions each of the most closely related genera *Pseudoxandra* and *Malmea*; two of Miliuseae; and a single representative of Piptostigmateae (*Annickia pilosa*) as most distant outgroup. For the *Mosannona* dataset, seven outgroup taxa were selected, exclusively from tribe Malmeeae: one accession each of *Cremastosperma*, *Ephedranthus*, *Klarobelia*, *Malmea*, and *Pseudomalmea*; and two of *Oxandra*.

Within *Cremastosperma*, 39 accessions included 24 of the 29 described plus two informally recognised species in Pirie et al. (2005b), from across the entire geographical distribution, with some species represented by multiple accessions. For the *Mosannona* dataset, 14 accessions included 11 of the 14 species recognised by Chatrou (1998). This compares to 13 samples/species of *Cremastosperma* and 7 of *Mosannona* represented in the analyses of Pirie et al. (2006).

### Character sampling

Character sampling differed somewhat between *Cremastosperma* and *Mosannona* matrices, depending on taxon-specific success with particular markers. DNA extraction, PCR and sequencing protocols for *Cremastosperma* followed (Pirie et al., 2005a, 2006), with modifications for *Mosannona* (as follows). For all 49 accessions of the *Cremastosperma* dataset the plastid encoded (cpDNA) markers *rbcL*, *matK*, *trnT-F* (at least partial) and *psbA-trnH* were sampled. Amplification and sequencing of a further cpDNA marker, *ndhF*, was only successful in 34 accessions and that of pseud*trnL-F* (an ancient paralogue of the plastid *trnL-F* region; Pirie et al., 2007), was only successful in 22 including just two outgroups (both species of *Malmea*).

For all 21 accessions of the *Mosannona* dataset, cpDNA *rbcL*, *matK*, *trnL-F*, *psbA*-*trnH* and the nuclear marker Phytochrome C were sampled; for 8 ingroup taxa and 3 outgroups additional cpDNA markers *atpB-rbcL* and *ndhF* and the nuclear marker malate synthase were also sampled. New primers were designed for phytochrome C, based on sequences of representatives of Magnoliales from GenBank (Magnoliaceae: *Magnolia* x *soulangeana*, Degeneriaceae: *Degeneria vitiensis*, Eupomatiaceae: *Eupomatia laurina*, Annonaceae: *Annona sp.*). Two forward and two reverse primers were designed: PHYC-1F: 5'-GGATTGCATTATCCGGC-3', PHYC-1R: 5'-CCAAGCAACCAGAACTGATT-3', PHYC-2F: 5'-CTCAGTACATGGCCAAYATGG-3' and PHYC-2R: 5'-GGATAGCCAGCTTCCA-3', applied in the combination 1F/2R (preferentially), 1F/1R or 2F/2R (where 1F/2R was not successful) and subsequently sequenced with 1F and 2R. Malate synthase was amplified in two overlapping pieces using the primers ms400F and ms943R (Lewis and Doyle, 2001), and mal-syn-R1 (5'-CATCTTGAGAAGATGATCGG-3') and mal-syn-F2 (5'-CCGATCATCTTCTCAAGATGATGTGG-3'), in the combinations ms400F/mal-syn-R1 and mal-syn-F2/ms943R. The thermocycler protocol for Phytochrome C followed that for *matK* (Pirie et al., 2005a); that for malate synthase was 94°C, 4min.; 35 cycles of (94°C, 1 min.; 59°C, 1 min.; 72°C, 2 min.); 72°C, 7 min.

### Sequence alignment and model testing

DNA sequences were edited in SeqMan 4.0 (DNAStar Inc., Madison, WI) and aligned manually. Gaps in the alignments were coded as present/absent characters where they could be coded unambiguously, following the simple gap coding principles of Simmons and Ochoterena (2000). Matrices are presented in Appendix 2 and on TreeBase (http://purl.org/phylo/treebase/phylows/study/TB2:S20848). We performed preliminary phylogenetic analyses of markers separately using PAUP* version 4.0 beta 10 (Swofford, 2003) (as below), to identify any differences between datasets, then individual markers were imported into SequenceMatrix (Vaidya et al., 2011) which was used to export concatenated matrices for further analyses. Best fitting data partitioning strategies (given models implemented in RAxML, MrBayes, and BEAST as below) were selected with PartitionFinder (Lanfear et al., 2012), given a concatenated matrix including all sequence markers and the 34 taxa for which *ndhF* was available, using a heuristic search strategy (‘greedy’) and comparison of fit by means of the Bayesian information criterion. Individual markers (each representing either coding or non-coding regions) were specified as potential data partitions.

### Phylogenetic analyses

Phylogeny was inferred under parsimony, using PAUP*; Maximum Likelihood (ML), using RAxML (Stamatakis, 2006); and Bayesian inference, using MrBayes version 3.2 (Ronquist et al., 2012). Under parsimony, heuristic searches of 1000 iterations, TBR, saving 50 trees per iteration were performed and bootstrap support was estimated for the markers individually and combined. Partitioned RAxML analyses of the nucleotide data only were performed including bootstrapping on CIPRES (Miller et al., 2010; Stamatakis et al., 2008). Bootstrapping was halted automatically following the majority-rule ‘autoMRE’ criterion. Two independent MrBayes runs of 10 million generations each were performed on the combined nucleotide and binary indel characters, implementing partitioned substitution models for the former, sampling every 1,000 generations. Convergence was assessed (using the PSRF) and post-burnin tree samples were summarised (using the sumt command) in MrBayes.

Given the phylogenetic results, we chose not to perform formal ancestral area analyses. The reasoning was first, that a realistic model for the biogeographic scenario would involve changing extents of areas through time, the definition of which would be likely to influence strongly the results. Second, the geographic structure in the phylogenetic trees (see Results) suggested a minimal number of range shifts, limiting the power of any parametric model (Pirie et al., 2012). We therefore adopt a parsimonious interpretation of the ancestral areas of the geographically restricted clades and use this to infer the timeframes for shifts in geographic range.

### Molecular dating

In order to infer the timing of lineage divergences within *Cremastosperma* and *Mosannona* we used BEAST 1.8.2 (Drummond et al., 2012) with a matrix of plastid markers (*rbcL*, *matK*, *trnL-trnF*, *psbA-trnH* and *ndhF*) of Malmeeae taxa combined from the individual matrices described above. Instead of using fossil evidence directly, that could only be employed across the family as a whole, we used two different secondary calibration points based on the Annonaceae-wide results of Pirie and Doyle (2012). Although results based on secondary calibration should be interpreted with caution (Graur and Martin, 2004; Schenk, 2016), we could thereby analyse a matrix including only Malmeeae sequences, avoiding the uncertainty and error associated with analysing relatively sparsely sampled outgroups with contrasting evolutionary rates. The original analyses were calibrated using the fossils *Endressinia* (Mohr and Bernardes-de-Oliveira, 2004), to constrain the most recent common ancestor (MRCA) of Magnoliaceae and Annonaceae to a minimum of 115 Mya, and *Futabanthus* (Takahashi et al., 2008), to constrain the MRCA Annonaceae to a minimum of 89 Mya. The latter fossil flower is incomplete and its membership of crown Annonaceae, although assumed in various studies, has not been tested with phylogenetic analysis (Massoni et al., 2015), but both of the two constraints individually imply similar ages for the clade (Pirie and Doyle, 2012). We used node age ranges derived using BEAST, assuming lognormal distribution of rates, and PL, assuming rate autocorrelation. Both of these assumptions are questionable in Annonaceae, and the methods result in somewhat differing ages. Under PL, Pirie and Doyle (2012) estimated the age of the Malmeeae crown node to be 52 Ma ± 3, which we represented with a) a uniform prior and b) a normal prior (mean of 52 and SD of 3) on the age of the root node. Under BEAST, they estimated 95% PP range of the age to be 33-22 Mya, which we represented with a) a uniform prior and b) a normal prior (mean 27, SD of 3). With these prior distributions we aimed to represent only uncertainty in the dating method (given the calibration), not the further uncertainty associated with fossil calibrations (i.e. that they represent minimum age constraints). We employed a birth-death speciation model, assumed a lognormal rate distribution, and rooted the tree by enforcing the monophyly of the two sister clades that together represent all the species, as identified in the previous analyses. We performed two independent runs of 10 million generations, assessed their convergence using TRACER version 1.6 (Rambaut et al., 2014), and summarised the results using programs of the BEAST package.

### Species distribution modelling

Data on occurrences of species included in the *Cremastosperma* and *Mosannona* phylogenies were extracted from the herbarium specimen database (BRAHMS; Filer, 2008) of Naturalis Biodiversity Center, Leiden, the Netherlands (http://herbarium.naturalis.nl/).

These comprised of records curated during our revisionary work on these genera based on specimens housed in multiple international herbaria. A total of 633 specimens of *Cremastosperma* and 442 specimens of *Mosannona* were available that were both identified to species by the authors and adequately georeferenced. These specimens included duplicates, i.e. multiple specimens of the same species collected either at the same locality, or not distantly enough from it to be treated as separate localities in our analyses. We only modelled species distributions for species recorded at least at four unique localities. After removal of duplicates the database consisted of 222 unique occurrence data points for 10 taxa of *Mosannona* and 319 data points for 20 taxa of *Cremastosperma*. The number of unique occurrence localities per species ranged from 4 (*Cremastosperma macrocarpum*) to 145 (*Mosannona depressa* subsp. *depressa*) (Appendix 3). Current environmental variables were downloaded from www.worldclim.org at 2.5 minute resolution and categorical FAO soil layers were downloaded from www.fao.org. In a preliminary analysis we used Pearson correlation tests to remove correlated climate (containing continuous data) and soil (containing numeric categorical data) variables, resulting in eight independent climate layers and ten independent soil variable layers (Appendix 4). We assessed all species distribution models (SDMs) using Maxent version 3.3.3k (Phillips et al., 2006). Maxent has been demonstrated to perform well when data are restricted to(1) solely presence only occurrences (Elith et al., 2006), and (2) small numbers of point localities (Wisz et al., 2008) (although the number of point localities needed is nevertheless higher for widespread species; van Proosdij et al., 2016). Given a set of point localities and environmental variables over geographical space, Maxent estimates the predicted distribution for each species. It does so by finding the distribution of maximum entropy (the distribution closest to uniform) under the constraint that the expected value of each environmental variable for the estimated distribution matches its empirical average over a sample of locations (Phillips et al., 2006). The resulting distribution model is a relative probability distribution over all grid cells in the geographical area of interest. It expresses the relative probability of the occurrence of a species in a grid cell as a function of the values of the environmental variables in that grid cell. Maxent runs were performed for each of the 29 taxa with the following options: auto features, random test percentage = 0, maximum iterations = 500.

A commonly used measure for model performance is the ‘area under the curve’ (AUC; Metz, 1978) that is calculated by Maxent. However, its utility has been criticized for reasons including erroneously indicating good model performance in the case of SDMs based on few records (Lobo et al., 2008; Raes and ter Steege, 2007). SDMs were tested for significance using a phylogenetically controlled null model approach (Raes and ter Steege, 2007). For each species, 100 replicate SDMs were generated using random locality points in the same numbers as the original locality points. The demarcation of the assemblage of locality points from which random samples are drawn is crucial, as a too large assemblage (e.g. the entire Neotropics) will almost certainly result in null model tests that indicate that SDMs are significantly different from random. We drew the random locality points from 1009 unique locality points where species of the tribe Malmeeae have been collected, which were extracted from the BRAHMS database of Naturalis Biodiversity Center, Leiden. *Cremastosperma* and *Mosannona* belong to this tribe that comprises ca 180 species (Chatrou et al., 2012). All of these species occur in wet tropical forests, and the majority have distributions that overlap with those of species of *Cremastosperma* and *Mosannona*. The replicate AUC values were sorted into ascending order and the value of the 95th element is considered as the estimate of the 95th percentile of the null model distribution. If the AUC value of the original SDM is greater than the 95th percentile it is considered significantly different from random.

Maxent output was used to calculate the pairwise Hellinger’s *I* niche overlap metric between all species within *Cremastosperma* and *Mosannona* using ENMtools (Warren et al., 2010). Hellinger’s *I* values were then used to calculate the average niche overlap within and between the different geographically restricted *Cremastosperma* and *Mosannona* clades. To test whether abiotic niches are phylogenetically conserved within both genera, we correlated pairwise Hellingers *I* values with pairwise patristic distances. These were calculated using PAUP*, using one of the most parsimonious trees, adding up connecting branches between species pairs under the likelihood criterion (GTR + gamma model, all parameters estimated). This approach allowed a meaningful interpretation of niche overlap, despite phylogenetic uncertainty within the geographically restricted clades that in most cases prevented the identification of sister-species pairs.

## Results

### Phylogenetic analyses

Variable and informative characters per DNA sequence marker are reported in Table 1. Per PCR amplicon, the plastid markers provided just 6-9/1-8 informative characters within *Cremastosperma*/*Mosannona*. Despite the short sequence length, pseud-*trnL-F* provided 13 within *Cremastosperma*.

**Table 1:**
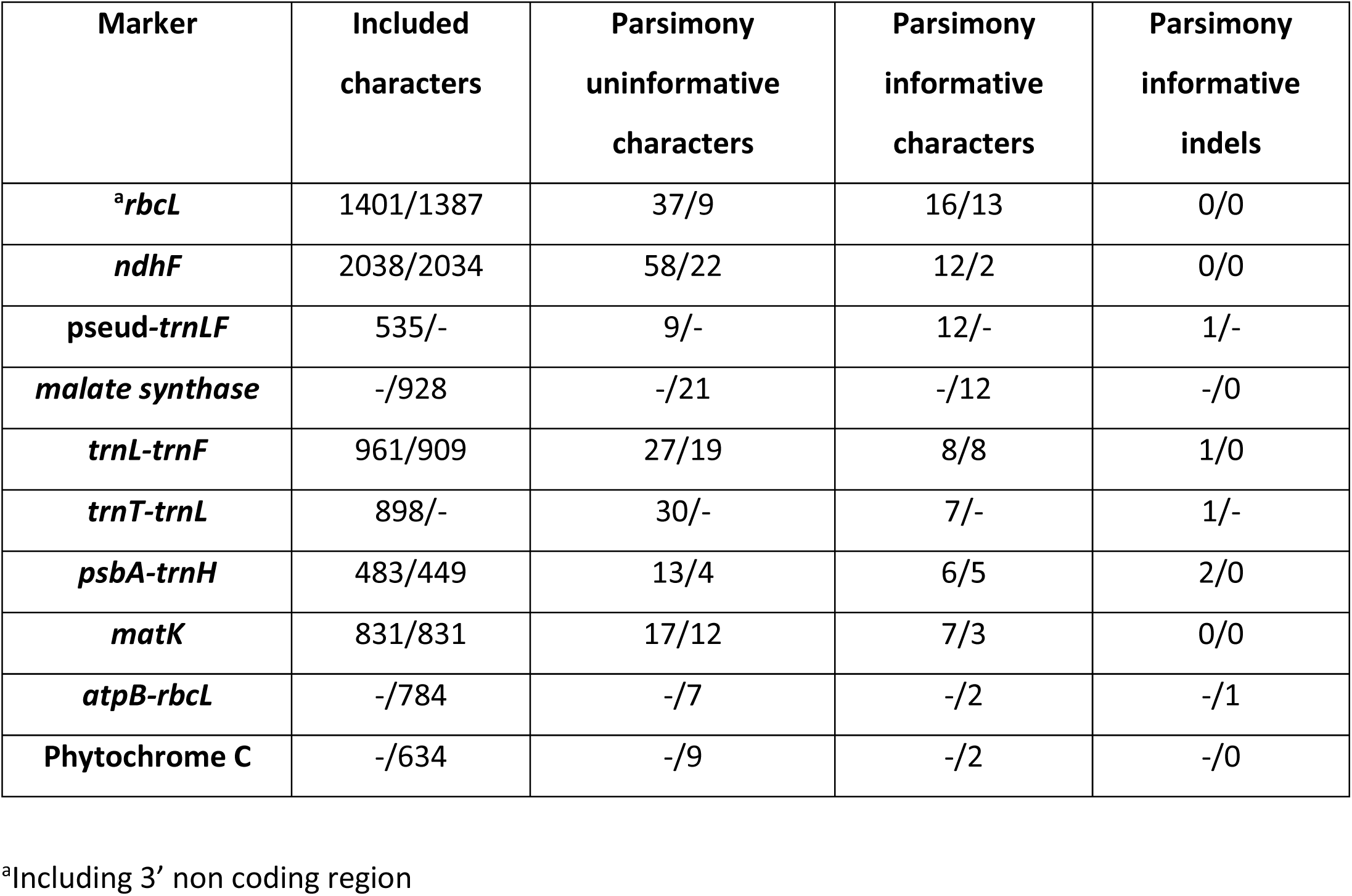
Included, variable and parsimony informative characters per DNA sequence marker, in descending order following the total number of informative characters in *Cremastosperma*. Values are made comparable by the inclusion of only *Cremastosperma*/*Mosannona* taxa for which all sequences were available. To compare variable/informative characters per sequence, note that *rbcL*, *ndhF* and malate synthase were PCR amplified in at least two fragments.

Parsimony bootstrap analysis of the individual data partitions revealed no supported incongruence (bootstrap support [BS] >70%). Data were thus combined in further analyses. Fig. 1 and Fig. 2 each show two best scoring trees as inferred under ML from the combined *Cremastosperma* and *Mosannona* data respectively. Fig. 1a and Fig. 2a show results including only taxa for which *ndhF* was available; and Fig. 1b and Fig. 2b show results including all taxa (i.e. with a considerable proportion of missing data). BS under parsimony and ML, and posterior probability (PP) clade support under Bayesian inference are also indicated. In both cases, the results of parsimony analyses were consistent with those obtained under ML and Bayesian inference, but analyses of the matrices with more missing data resulted in a greater number of nodes subject to PP≥0.95 than those subject to BS≥70.

**Fig. 1:**
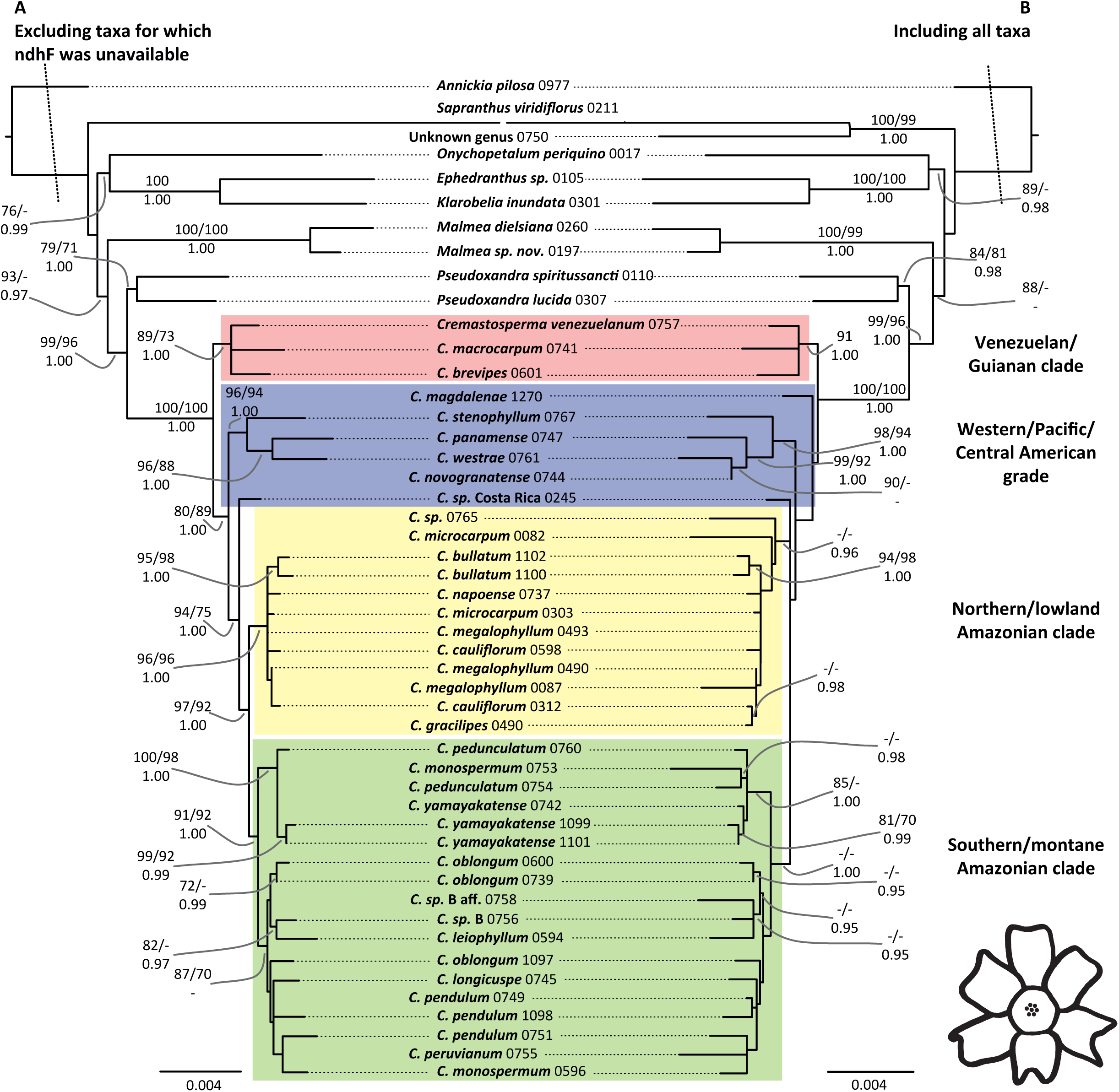
Phylogenetic hypotheses for *Cremastosperma* based on *rbcL*, *matK*, *trnT-F*, *psbA-trnH*, *ndhF*, and pseud*trnL-F*. A: Excluding taxa for which *ndhF* was unavailable; B: including all taxa. Topologies and branch lengths are of the best scoring ML trees with scale in substitutions per site. Branch lengths subtending the ingroup are not to scale. Clade support is indicated: ML and parsimony bootstrap percentages (above; left and right respectively) and Bayesian posterior probabilities (below); as are major clades referred to in the text.

**Fig. 2:**
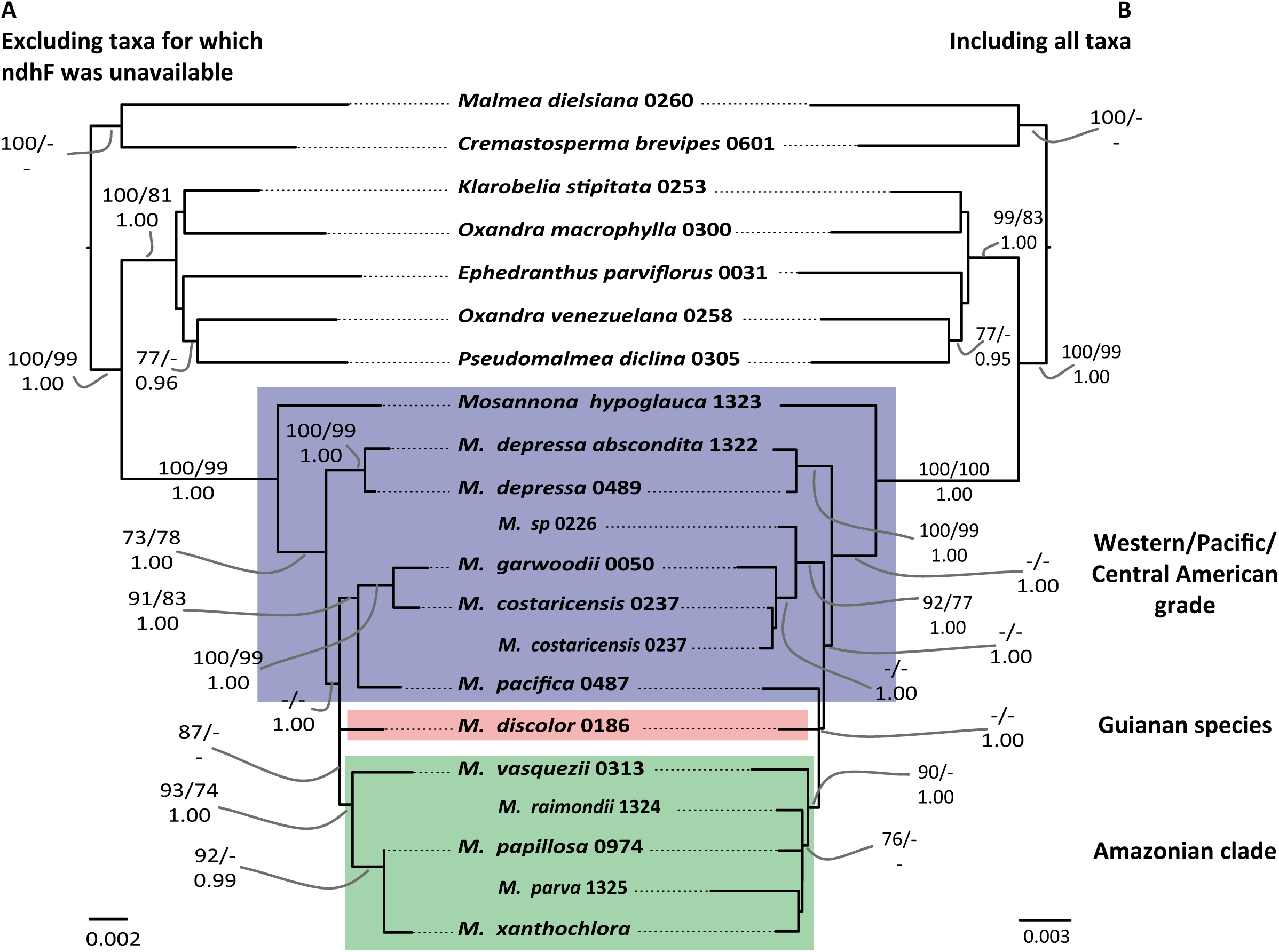
Phylogenetic hypotheses for *Mosannona* based on *rbcL*, *matK*, *trnL-F*, *psbA-trnH*, *ndhF*, *atpB-rbcL*, PHYC, and malate synthase. A: Excluding taxa for which *ndhF* was unavailable; B: including all taxa. Topologies and branch lengths are of the best scoring ML trees with scale in substitutions per site. Branch lengths subtending the ingroup are not to scale. Clade support is indicated: ML and parsimony bootstrap percentages (above; left and right respectively) and Bayesian posterior probabilities (below); as are major clades referred to in the text.

The combined analyses revealed a number of clades within both *Cremastosperma* and *Mosannona* corresponding to discrete geographic areas. In *Cremastosperma*, the divergence of the Venezuelan and Guianan lineages, together in a well-supported clade, occurred prior to that leading to clades found in the tropical Andes or in the Chocó/Darién/western Ecuador region or Central America (i.e. either the west or the east side of the Andes mountain chain). Western species fall within a grade of three lineages (a clade including *Cremastosperma panamense*, *C. novogranatense*, *C. stenophyllum* and *C. westrae*; and two isolated lineages corresponding to *C. sp*. A from Costa Rica and *C. magdalenae* from the Magdalena valley of Colombia), in which the three Central American species are not each other’s closest relatives. A single clade including all Amazonian species is nested within this grade; it comprises two clades: one including more lowland and northerly distributed species; the other more southerly and higher elevation species. In *Mosannona* the species are similarly represented by a western grade and a single eastern, Amazonian, clade. The single Guianan species (none are known from Venezuela), rather than being sister to these, is nested within, part of a polytomy of western and the eastern lineages.

### Molecular dating

Relaxed-clock molecular dating using BEAST resulted in ultrametric trees with topologies consistent with those obtained separately for *Cremastosperma* and *Mosannona* under parsimony, ML, and Bayesian inference (Fig. 3). The posterior age distributions for the root node given the normal prior were somewhat wider than the priors (37-21 Mya instead of 33-22 for the BEAST calibration; 69-47 instead of 52 +/- 3 for the PL calibration), reflected in somewhat wider/older age estimates for shallower nodes. The nodes that define the divergence of lineages to the west and east of the Andes are A) the crown nodes of the clades including exclusively Western species (such as *M. pacifica*, *M. garwoodii*, *M. costaricensis* and *M. sp*.: Fig. 3 A1; and *C. stenophyllum*, *C. panamense*, *C. novogranatense* and *C. westrae*: Fig. 3, A2); B) the stem nodes of the exclusively Amazonian clades (Fig. 3, B1 and B2), representing the age of the most recent common ancestors of western and eastern lineages; and C) the crown nodes of Amazonian clades (Fig. 3, C1 and C2). The minimum and maximum ages of dispersal into Central American are defined by the crown and stem nodes respectively of Central American lineages. In the case of *Cremastosperma* these are single species/accessions and hence without crown nodes in these phylogenetic trees. In the case of *Mosannona*, two clades of taxa are endemic to Central America (*M. depressa* sspp., and *M. costaricensis*/*M. garwoodii*) for which both stem and crown node ages could be estimated. The age ranges for these nodes given the older and more recent secondary calibrations (root node 25-13 Ma; 12-3 Ma) are reported in Table 2. The confidence intervals for each of A1 and A2, B1 and B2 and C1 and C2 for a given calibration largely overlap. The implied time window for a putative vicariance process caused by Andean uplift ranges between 24 Ma and 7 Ma (older calibration) and 13 Ma and 3 Ma (recent calibration). The minimum age for dispersal into Central America ranged from 15 to 1 Ma, and only given the older calibration did the minimum estimate for the *M. costaricensis*/*M. garwoodii* crown node exceed 3.5 Ma (5 Ma).

**Fig. 3:**
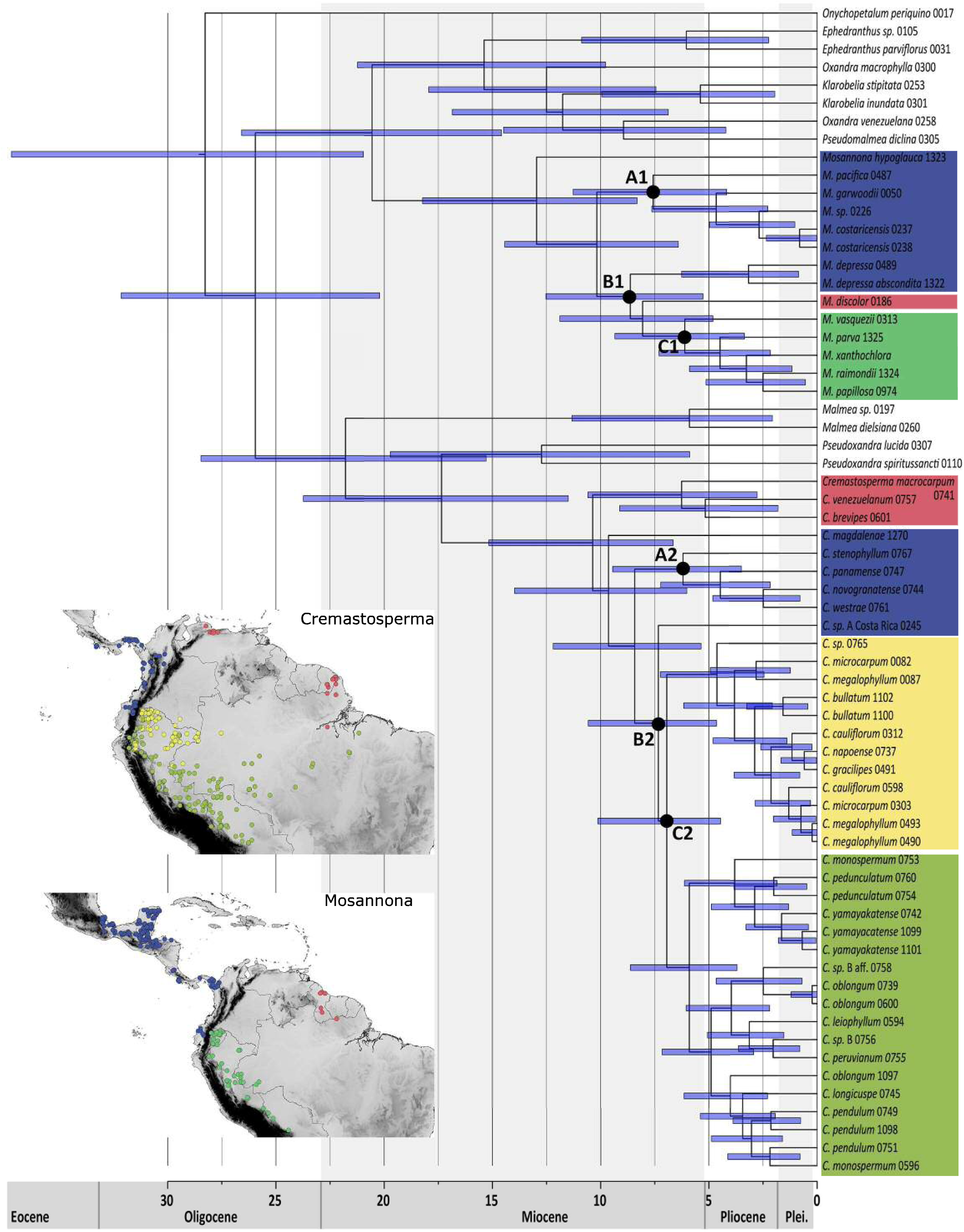
Chronogram of Malmeeae (given the normal distributed age calibration), indicating nodes within *Cremastosperma* and *Mosannona* that define the divergence of lineages to the west and east of the Andes, as referred to in the text and in Table 2. Distributions of the clades, plus those of the *Cremastosperma* northern/lowland and southern/montane clades are illustrated on the inset maps.

**Table 2:**
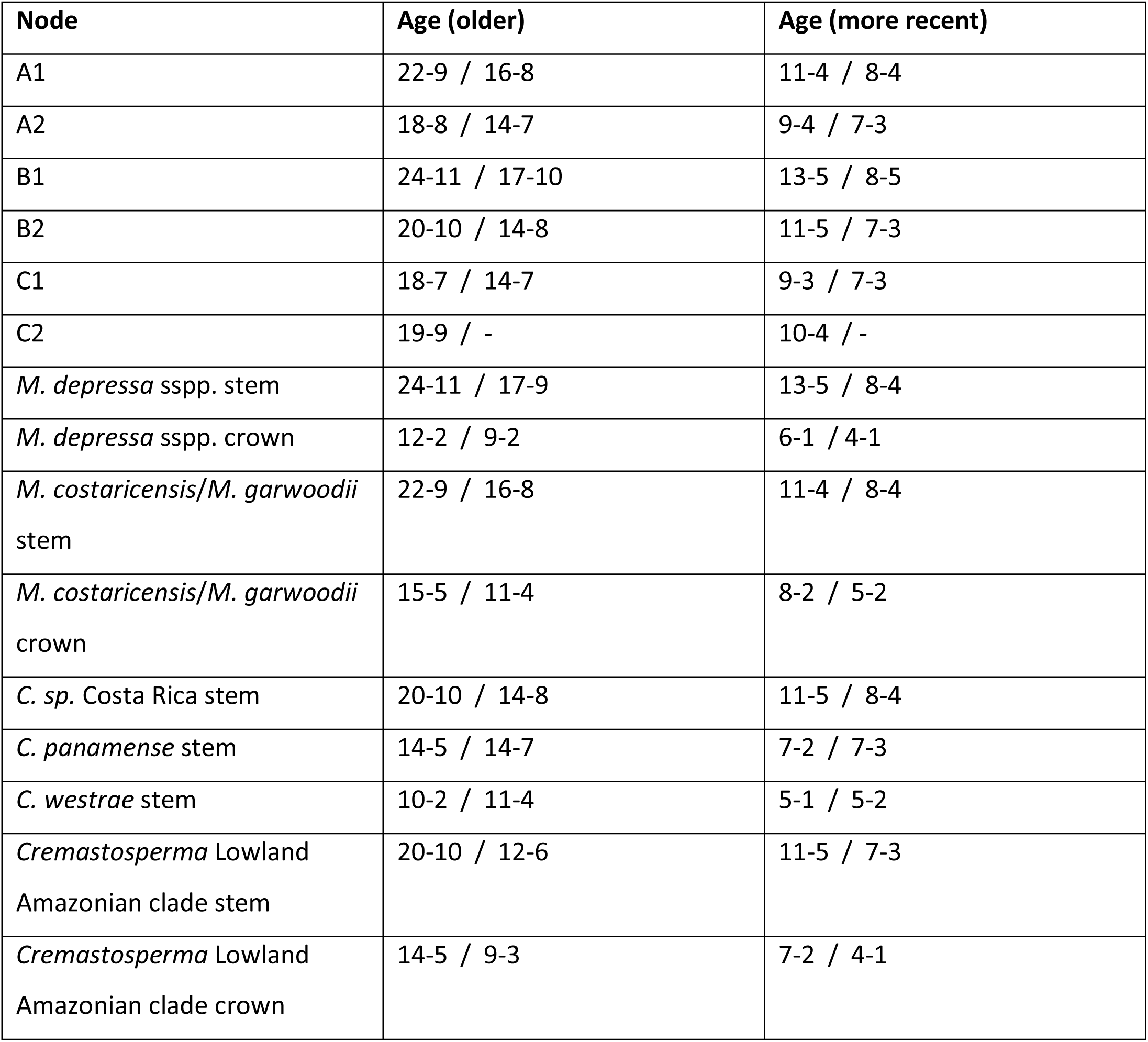
Relaxed clock molecular dating estimates (limits of the 95% HPD) for the ages of selected nodes given older and more recent secondary calibration points. Ages before the forward slash result from analyses with normal distributions of the age prior, after the forward slash result from analyses with uniform distributions of the age prior.

### Species distribution modelling

Null-model tests showed that all species distribution models were significantly different from random, except for that of *Cremastosperma pendulum,* which was excluded from subsequent analyses. Pairwise intra-generic Hellingers *I* niche overlap measures ranged from 0.009 (*M. depressa* subsp. *depressa* vs. *M. xanthochlora*) to 0.987 (*C. pacificum* vs *C. novogranatense*). Niche similarity was found to be negatively correlated with phylogenetic distance in *Cremastosperma* (squared Pearson correlation R^2^ = 0.1933, *p* < 0.0001), while it was not in in *Mosannona* (squared Pearson correlation R^2^ = 0.0392, *p* = 0.2207).

Niche similarity is considerably greater for Amazonian species than for species in any of the remaining areas (Table 3). In contrast, the smallest niche overlap for species within a given area was observed for the areas west of the Andes, viz. Pacific South America and Central America. Niche similarity within each of the four areas exceeds niche similarity between them in all cases. Only the mean value for Hellinger’s *I* within Pacific / Central America approaches those for between-area niche similarity. For *Cremastosperma*, the two areas with the largest niche differences are the Amazonian lowland and the Guianas / Venezuela. For *Mosannona* niche overlap cannot be calculated for the Guianas and Venezuela due to the occurrence of only single species, yet the within-area niche differences are larger in the Amazonian lowland than Pacific South America and Central America, as in *Cremastosperma*.

**Table 3:**
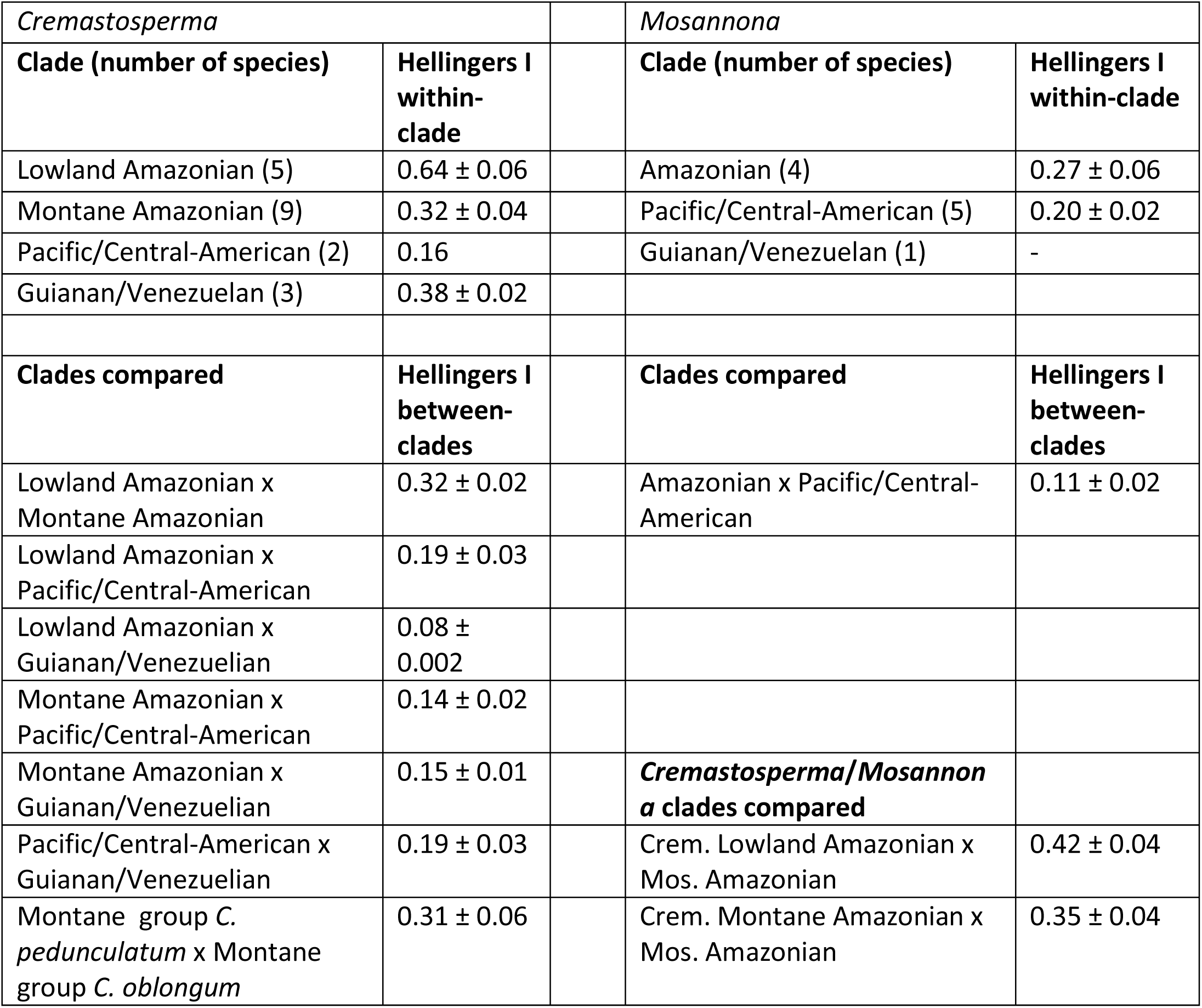
Within and between-area comparison of niche overlap (Hellinger’s *I*).

## Discussion

### Consistent biogeographic signals revealed in phylogenies of two rainforest tree genera

The phylogenetic hypotheses for *Cremastosperma* and *Mosannona* show strong geographic signal, with clades endemic both to regions divided by obvious present day dispersal barriers (east and west of the Andes), and to regions that are currently largely contiguous (i.e. within Amazonia). These results appear to contrast with the assumptions of rapid dispersal abilities of Amazon biota (Dick et al., 2003; Pennington and Dick, 2011), such as species-rich Neotropical lowland clades (Dexter et al., 2017) including the more typically Amazon-centred, more widely distributed, Annonaceae genus *Guatteria* (Erkens et al., 2007). The strong correlation between phylogenetic patterns and geographic areas found here resembles more closely the patterns found in Neotropical birds (Smith et al., 2014).

Andean-centred distribution patterns are the exception in Neotropical Annonaceae clades, but have originated multiple times independently, for example in *Malmea*, *Klarobelia*, and *Cymbopetalum*, as well as *Cremastosperma* and *Mosannona* (Pirie et al., 2006). This may reflect an ecological distinction between more montane Andean-centred genera and their widespread lowland sister-groups. It may also be associated with more limited dispersal abilities of the smaller understorey trees typical of *Cremastosperma* and *Mosannona*, as opposed to canopy trees (Kristiansen et al., 2012) as represented e.g. by species of *Guatteria*. Similarities between different Andean-centred clades suggests the potential for identifying common and more general underlying causes.

### The impact of the Andean orogeny on lowland rainforest taxa in the Neotropics

The diversifications of extant species of *Cremastosperma* and *Mosannona* date to the Middle-Late Miocene (Fig. 3). During this period, the uplift of the westernmost Cordillera Occidental, the oldest of three cordilleras in Colombia, and of the Cordillera Central had been completed (Fig. 5). The easternmost Cordillera Oriental was the last Colombian cordillera to rise, with uplift starting around 13-12 Ma. The Cordillera Oriental is decisive for the establishment of a dispersal barrier for lowland organisms on either side of the Andes. During the Miocene it consisted of low hills, only attaining its current height during the Pliocene and early Pleistocene (Gregory-Wodzicki, 2000), and as a result the northwestern part of the current Amazon basin was contiguous with areas now in the Colombian states of Chocó and Antioquia, west of the Andes.

Rough bounds for the timing of establishment of an east-west dispersal barrier can be inferred from stem and crown node ages of exclusively eastern and western clades. In both *Cremastosperma* and *Mosannona* the stem node ages, representing common ancestors that were either widespread or which were able to disperse between areas, date to 24-10 Ma (older estimates) or 13-5 Ma (more recent estimates). Crown node ages, representing common ancestors after which there is no further evidence for such dispersal, date to 22-7 Ma (older estimates) or 11-3 Ma (more recent estimates). The error margins are wide, but consistent with a vicariance scenario. Comparable evidence for the Andes forming a strong barrier to dispersal have been found in phylogeographic patterns of individual species (Dick et al., 2003; Scotti-Saintagne et al., 2013), and phylogenies of clades of lowland tree and palm species (Antonelli et al., 2009; Barfod et al., 2010; Roncal et al., 2013). However, a notable contrast is presented by results in tropical orchids, for which there is evidence of more recent trans-Andean dispersal, suggesting that the barrier is less effective for plants with more easily dispersed diaspores (Pérez-Escobar et al., 2017a, 2017b).

### The drainage of lake Pebas, dispersal and diversification in western Amazonia

prior to the Andes forming a dispersal barrier for lowland rainforest biota, dispersal from northwestern South America further into the Amazon may have been blocked by lake Pebas, that flooded the entire western Amazon from c. 17 to c. 11 Ma (Hoorn et al., 2010; Wesselingh et al., 2002). One of the first areas to emerge in the upper Amazon basin was the Vaupes Swell or Vaupes Arch, which may have acted as a dispersal route from western Amazonia to the Guianas for frogs during the Late Miocene (Lötters et al., 2010). This route is also plausible in the case of *Cremastosperma*. The divergence of the Venezuelan/Guianan clade was prior to that of western and eastern clades, which suggests either an earlier widespread distribution, or stepping stone dispersal, across northern South America. The absence of *Mosannona* in rainforest of coastal Venezuela could be explained by extinction or failure to sample species (the taxa known to this region are rare; Chatrou and Pirie, 2005). However, the differences between the two genera both in relationships and in the niches of the Guianan species compared to other clades may instead point to different origins. Whilst *Cremastosperma* Venezuelan/Guianan clades are both distantly related and occupy more dissimilar niches compared to Amazonian clades, the Guianan *M. discolor* is closely related to the *Mosannona* Amazonian clade and similar in niche. This may imply a more southerly dispersal route that could have occurred much more recently.

Despite the broad similarities between *Cremastosperma* and *Mosannona*, the clades also differ in species numbers, particularly reflected in fewer Amazonian taxa in *Mosannona*. In *Cremastosperma*, we identified a single Amazonian clade comprising two subclades of contrasting distributions: one of more lowland, northerly species, the other of more montane or southerly distributed species (Fig. 1; Fig. 3). These patterns reflect clade-specific niche differences, whereby species of the lowland/northern clade occupy a particularly narrow niche compared to the montane/southerly clade.

Given its distribution and age (crown node between 14 and 2 Ma old), the lowland/northern clade may provide further evidence of diversification within the forested habitat that formed following drainage of Lake Pebas. A similar result was obtained within the palm genus *Astrocaryum* (Roncal et al., 2013), whilst in Quiinoideae diversification was earlier, with subsequent independent colonisations of the region (Schneider and Zizka, 2017). The distributions of the individual species of lowland/northern *Cremastosperma* broadly overlap, but they are morphologically distinct and particularly diverse in floral characteristics, with large differences in flower colour, indumenta, and inflorescence structure. This might point to sympatric speciation in the group, perhaps driven by pollinators or herbivores, as demonstrated for some Neotropical clades of woody plants (Alcantara et al., 2014; Kursar et al., 2009).

The montane/southern *Cremastosperma* clade by contrast is morphologically more homogenous, spread over a wider latitudinal range and with lesser overlap in the distributions of individual species (with the exception of the widespread *C. monospermum*, which alone represents much of the northern and eastern extremities of the clade; Fig. 3). Given the age of this clade and its proximity to the Andes, diversification may have been influenced by habitat shifts caused by uplift and/or Pleistocene climatic fluctuations. The large proportion of such pre-montane species across the *Cremastosperma* clade as a whole may explain the lack of niche-based evidence for allopatric speciation, with the significant negative correlation between phylogenetic distance and niche overlap (Fig. 4A) reflecting (phylogenetically conserved) niche differentiation. Both this scenario and that for the lowland/northern clade differ markedly from that of allopatric speciation with lack of niche shifts inferred in the diversification of lowland *Monodora* and *Isolona* (Annonaceae) in Africa (Couvreur et al., 2011).

**Fig. 4:**
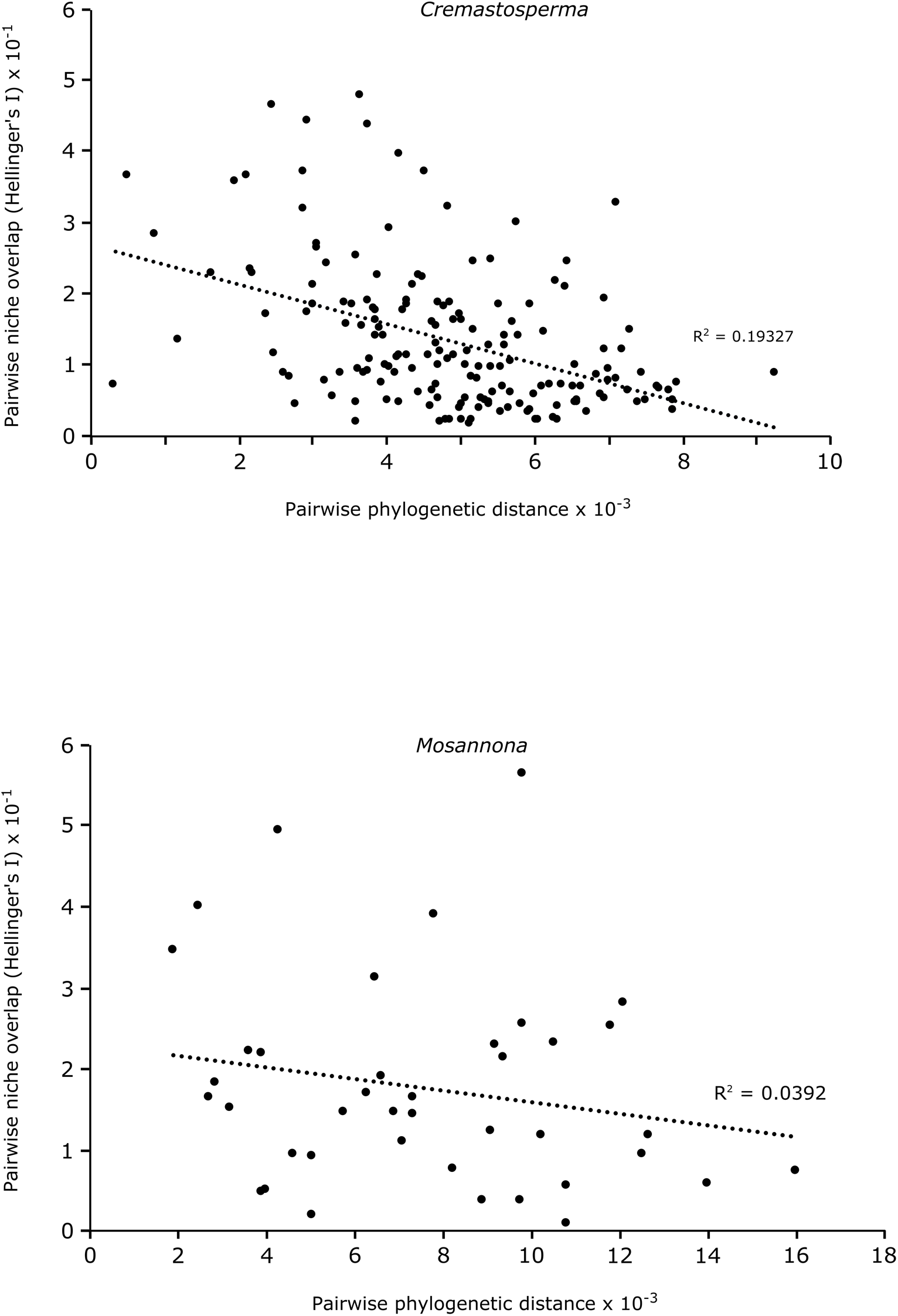
Graph showing the correlation per genus between Hellinger’s *I* metric and patristic distance.

In *Mosannona*, the single Amazonian clade represents far fewer species and ecological distinctions between clades are not so obvious. The overall niche overlap is lower than that of the *Cremastosperma* lowland Amazon clade, suggesting that *Mosannona* may have failed to adapt to the same habitat and, perhaps as a result, failed to diversify to the same extent. Neither niche conservatism, nor a negative correlation between phylogenetic distance and niche overlap as seen in *Cremastosperma*, are significant across the genus as a whole. This may in part reflect the lower numbers of species and/or records available for analysis. Moreover, the use of remotely sensed climate data could have improved the predictive power of our species distribution models (Deblauwe et al., 2016). Nevertheless, despite the lack of evidence from ecological niches and given the entirely non-overlapping species distributions, allopatric speciation in this and the wider clade is still plausible.

### Closure of the Panama Isthmus and geodispersal into Central America

Numerous studies have assumed an age of 3.5 Ma from which dispersal between North-Central and South America would have been possible. However, evidence has been presented suggesting that plant dispersal across the Panama isthmus occurred earlier than that of animals (Cody et al., 2010) and that dispersal occurred prior to its complete closure (Bacon et al., 2015). Winston et al. (2017) demonstrated that strictly terrestrial South American army ants colonized Central America in two waves of dispersal, the earliest of which occurred 7 Ma. However, pre-Pliocene dates for the formation of an isthmus are still disputed (O’Dea et al., 2016), and further data is warranted to further test the timing of dispersal events, potentially between separate land masses, in different groups.

Of the species of *Cremastosperma* and *Mosanonna* that are found in Central America, only a few extend beyond Panama or Costa Rica and some, such as *C. panamense* and *M. hypoglauca*, straddle the Panama/Colombian border with populations in both Central and South America. Central American lineages of *Cremastosperma* are single species nested within western clades, implying multiple independent northerly dispersals. The stem lineages date to 20-1 MY, and these dispersal events must have taken place subsequently, but given the data available it is not possible to further narrow down the timeframe. *Mosannona* include two clades of exclusively Central American species (*M. garwoodii* and *M. costaricensis*) or subspecies (*M. depressa*), also nested within a western grade, and the ages of these crown groups can be interpreted as minima for the preceding northerly dispersals. Although independent arrivals over earlier land connections (Bacon et al., 2015) or even (long-distance) dispersal (Hughes et al., 2013; Pennington and Dick, 2011) cannot be ruled out, the ages of these nodes, ranging from 12-1 and 15-2 Ma would not exclude a scenario of concerted range expansions (i.e. geodispersal) following Pliocene closure of the isthmus.

The biogeographic scenario that emerges from our study reinforces the importance of the geological history of north-western South America during the Late Miocene − Early Pliocene for the evolution of plant diversity in the Neotropics. Three major geological processes (the uplift of the Andes, lake Pebas, and the Panama isthmus) interacted within a timeframe sufficiently narrow to challenge the discerning power of current molecular dating techniques. Ancestral lineages that were present in north-western South America were subject to vicariance and to opportunities for dispersal within a period of little more than about 4 million years. These events nevertheless left common signatures in the phylogenies of clades such as *Cremastosperma* and *Mosannona* (Fig. 5) that in part − though not entirely − are reflected in the similarities in their modern distributions.

**Fig. 5:**
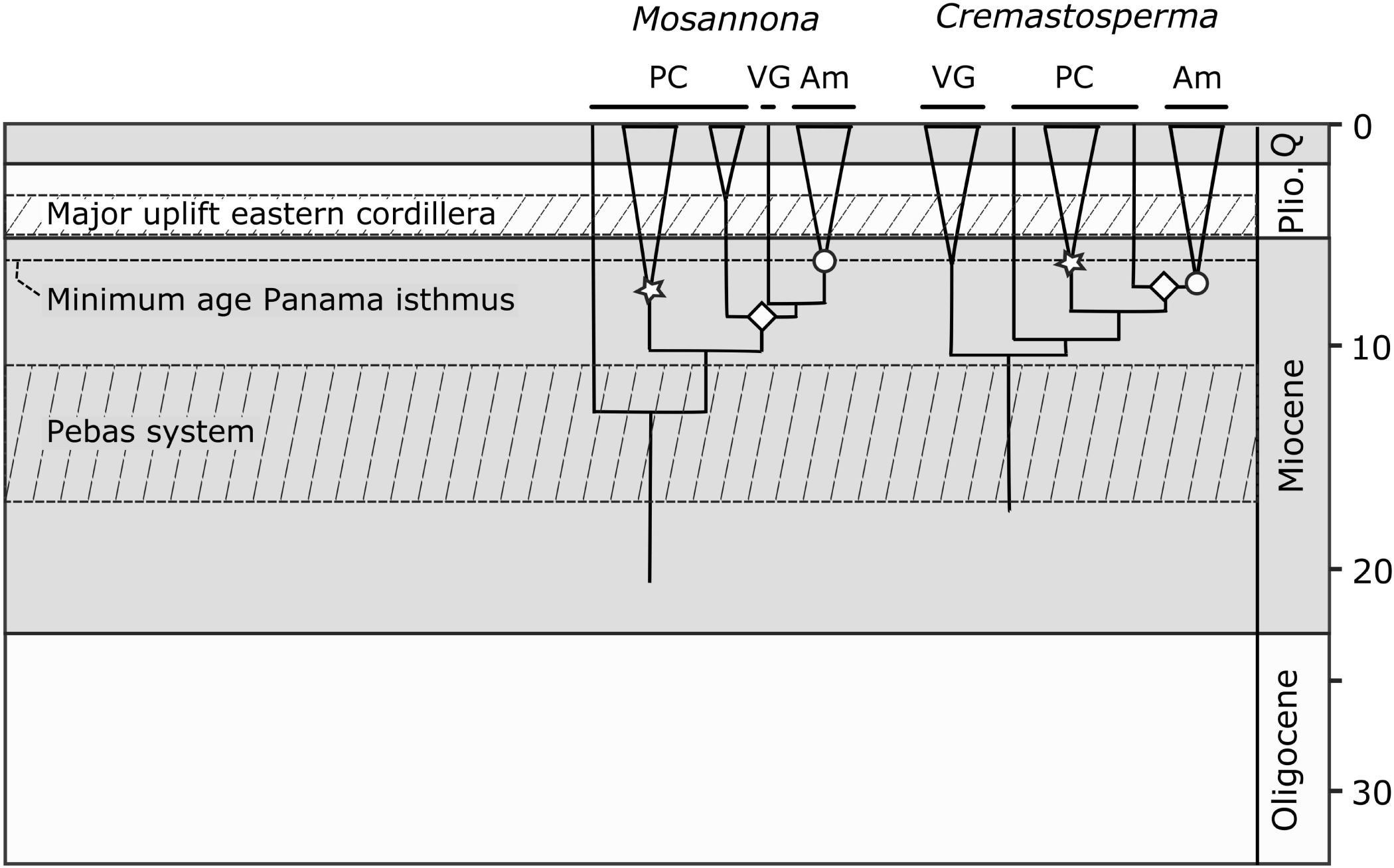
Scheme of geological events and ages of clades in *Cremastosperma* and *Mosannona*.

## Acknowledgments

The authors acknowledge The Netherlands Organisation for Scientific Research (NWO; R 85-351 to MDP) and the Hugo de Vries Foundation (to LWC) for support of fieldwork, and Jan Wieringa for assistance with the BRAHMS database. Constructive criticism from Hervé Sauquet and Thomas Couvreur as reviewers/recommenders for PCI Evol Biol (https://evolbiol.peercommunityin.org) on a earlier draft of the paper is gratefully acknowledged.

## Appendix 1

Accessions details

## Appendix 2

Sequence matrices

## Appendix 3

Numbers of unique records and area under the curve per species included in the species distribution modelling.

## Appendix 4

The eight independent climate layers and ten independent soil variable layers used for species distribution modelling.

